# Metrics for Distinguishing Biological Versus Interventional Change in AI Models

**DOI:** 10.64898/2026.06.24.733252

**Authors:** M. Antony Ewing

## Abstract

Statistical and machine-learning models of longitudinal biological data evaluate change by comparing each new observation against the trajectory implied by prior observations, assuming the process generating that trajectory is stable. We use “data substrate” to mean the underlying structure of the longitudinal data that determines what any such model can recover, independent of its architecture or capacity. When the generating process changes — whether through a biological transition or through an external intervention — the prior trajectory ceases to be a valid reference, and extrapolated predictions can be confidently wrong with no internal signal that the reference has failed. A distinct and recognised difficulty is that biological change and interventional change, observed only through serial intertemporal comparison under an assumed trajectory, are readily conflated; existing approaches address this through causal assumptions or hidden-confounder models rather than from the data substrate itself. Here we ask whether the two can be distinguished at the substrate level, and we introduce two subject-level metrics that quantify the geometric signature an interventional change leaves in the data: Curvature Shift, the change in trajectory slope across the event, and Deformation Risk, the departure of post-event observations from the prior-trajectory reference. We evaluate the condition on longitudinal cognitive measurements from 309 human subjects in the Alzheimer’s Disease Neuroimaging Initiative (ADNI), a large longitudinal dataset containing two distinct, ex-ante-defined regime-change events in the same subjects: a biological transition and an intervention. A model extrapolating the pre-event trajectory assigned the wrong direction of change to roughly two-thirds of post-event observations (post-event sign accuracy 0.341 after the biological event and 0.350 after the intervention, against a chance value of 0.50); only 11% of postbiological-event and 12% of post-intervention readings remained concordant with prior dynamics, and a higher-capacity multilayer perceptron reproduced rather than resolved the error. Curvature Shift was 2.23-fold higher after the biological event (p = 4.4×10-8) and 2.26-fold higher after the intervention (p = 7.4×10-8), and the two metrics were coupled (ρ= 0.500; 95% CI, 0.407–0.587). Findings replicated on an independent endpoint and survived propensity matching, permutation, 1 and leave-one-out. The metrics detect, per subject, when a fitted model’s reference has stopped governing the data and whether the departure carries the geometric signature of an interventional change.

## Introduction

This paper is a companion to an earlier study6 in which we considered how data substrates can be evaluated for adequacy in supporting AI biological-model predictions when model weights and internals are unavailable. In that paper it was established that AI models are generally suitable for determining state-to-state predictions but struggle with state-to-changing-state predictions when the data are not encoded to include changing-state dynamics. As a result, an assessment of the data substrate can be performed that partitions the data into encoded and nonidentifiable components, where only the former admit state-to-changing-state predictions. Here we extend this analysis by offering two additional metrics for distinguishing changing-state dynamics that arise from different processes: biological change and interventional change. Under biological change, an organism is observed to follow a natural change in its course trajectory that may appear to be a response to an external intervention when only serial, intertemporal comparisons are made under the assumption of a certain trajectory. In that case, biological and interventional change are conflated. As noted in the companion paper, when a model — particularly a large language model — is complex and opaque, with weights and benchmark substrate unknown to the user or to observers, such conflation is a significant challenge. However, by again examining the data substrate, biological and interventional change can be distinguished using additional metrics that determine whether such distinctions are possible and under what circumstances. In particular, we show that interventional change, by definition, requires a geometric change that possesses a detectable signature in a given data substrate. The difficulty this paper addresses is not new. Distinguishing the natural course of a biological process from the effect of an intervention is a long-recognised and active problem in the analysis of longitudinal biological data. The two are known to leave geometrically distinct patterns: a process that merely shifts the level of a trajectory while preserving its rate is qualitatively different from one that alters the rate itself, and a change in the rate of progression is the recognised hallmark of a process-modifying effect 3. Yet recovering this distinction from observational data alone is difficult.

**Figure 1.**
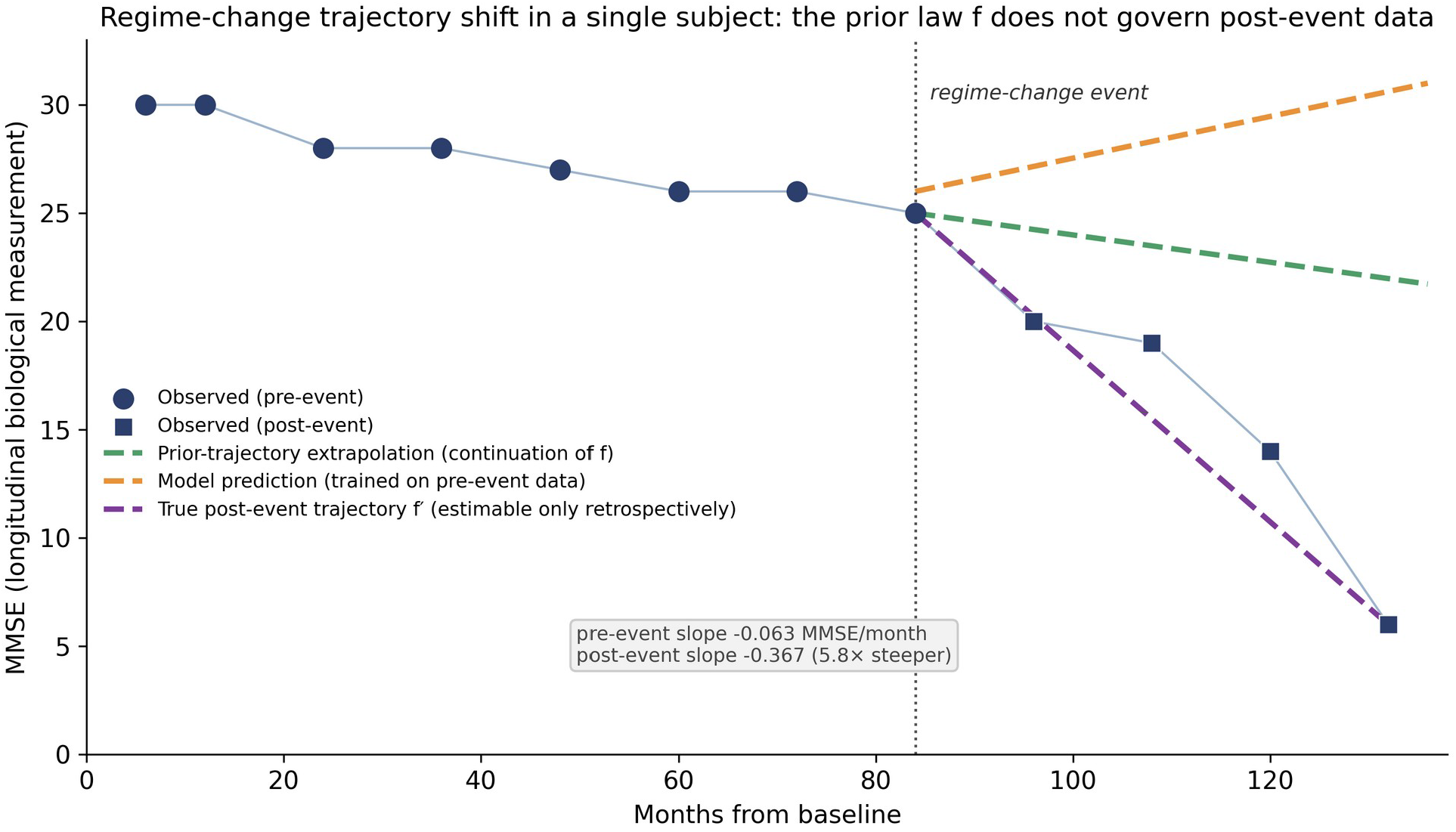
Regime-change trajectory shift in a single subject. The prior law f governs the pre-event observations; after the regime-change event the data are generated by a shifted law f′. Both extrapolation of the prior trajectory and a model trained on pre-event data miss the post-event acceleration, which is recoverable only retrospectively. The post-event slope is roughly six times steeper than the pre-event slope.

## Methods

### Data substrate and cohort construction

We used longitudinal measurements from ADNI, a large longitudinal dataset of human biological subjects. A single brief repeated measure recorded at nearly every visit (MMSE) served as the primary longitudinal signal; an independent endpoint (CDR Sum-of-Boxes) was used for replication. The analysis is defined entirely on the observed measurement sequence and its rate of change; no model weights, activations, or training data enter any quantity below.

For each subject we ordered all visits by month and formed consecutive inter-visit transitions. For visit *i* with measurement value x_i at month t_i, the transition to visit *i*+1 has elapsed time Δt = t_{i+1} − t_i (transitions with Δt ≤ 0 were discarded) and per-month rate of change

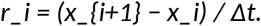

The per-month rate r (denoted delta_y_per_month) is the unit of analysis throughout; the running measurement value x_i is the covariate against which rate is modeled.

Subjects were retained when they had at least MIN_TOTAL_VISITS recorded measurements and at least two transitions in each phase (MIN_PRE_VISITS = 2, MIN_POST_VISITS = 2). Applying these filters yielded a primary cohort of 309 subjects (97 transition, 212 stable) for the biological-event arm, and a parallel intervention arm of 59 subjects with 204 stable controls.

### The two ex-ante regime-change events and phase assignment

Two regime-change events were defined ex ante in the same subjects, each splitting a subject’s transitions into a pre-event and a post-event phase.

For the **biological event**, a subject was labeled a transition subject if the dataset’s recorded state reached the terminal biological category at any visit; the event month was the month of the first such visit. A transition with start month t_i was assigned to the pre phase if t_i < event month and to the post phase otherwise.

For the **interventional event**, the event month was the first recorded onset of the external intervention, and transitions were split into pre and post phases by the same rule.

For **stable subjects**, in whom no regime change occurs, the phase boundary was set to the subject’s median visit month, so that the identical two-phase pre/post construction could be applied as a negative control.

### Per-subject trajectory fits

Within each phase, the per-month rate was modeled as a linear function of the running measurement value by ordinary least squares. For the pre-event phase this yields a slope b_pre and intercept a_pre; for the post-event phase, b_post and a_post. A fit was computed only when the phase contained at least two finite observations with non-degenerate spread in the covariate (standard deviation > 1×10^−12^); subjects lacking a valid fit in either phase were excluded. The pre-event fit defines the **prior trajectory** (the prior law), used as the reference against which post-event observations are tested:

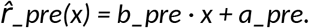

### Substrate-level metrics

Two subject-level metrics are computed from the subject’s own measurement record. Both are properties of the data substrate: they require no model weights, counterfactuals, or assumed causal model, and apply identically whether the underlying predictor is a linear fit, a recurrent network, or an opaque black-box system.

**Curvature Shift (CS)** is the absolute change in the rate-vs-state slope across the event:

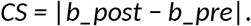

*CS near zero indicates that the generating rate is preserved across the event; a large CS indicates a change in the generating rate — the geometric signature of an interventional or process-modifying change. CS is reported in measurement units per month.*

**Deformation Risk (DR)** is the mean absolute departure of the post-event observations from the prior-trajectory reference. For the n_post post-event observations with covariate values x_j and observed rates r_j,

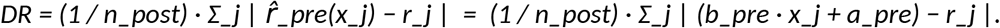

*DR measures the degree to which the prior law has ceased to govern the subject. As a within-subject reference scale, the analogous pre-event residual (the same quantity evaluated on the pre-event observations against their own fit) is reported, and the post-event-to-pre-event ratio of these residuals summarizes the inflation in departure across the event.*

### Direction accuracy and the four post-event categories

For each post-event observation, the **sign accuracy** of the prior trajectory is the fraction of observations whose realized direction of change matches the direction predicted by the prior-trajectory reference, sign 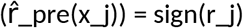; a value of 0.50 corresponds to chance. The pre-event sign accuracy is the same quantity evaluated within the pre-event phase as an internal reference.

Each post-event observation was assigned to one of four mutually exclusive categories by comparing what the prior law predicts 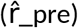, what the post-event (deformed) law predicts 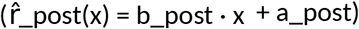, and what the prior-law residual implies:

- **concordant** — the prior-law and post-event-law predictions agree in direction 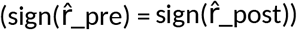 and the observation is consistent with the prior trajectory;
- **silent shift** — the prior-law and post-event-law predictions disagree in direction 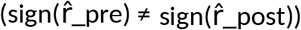 while the observed direction still superficially tracks decline, so that the rate no longer belongs to the prior law though direction alone does not reveal it;
- **apparent decline** — the prior-law residual reads as worse-than-expected while the post-event law is non-declining;
- **apparent improvement** — the prior-law residual reads as better-than-expected while the post-event law is declining.

In stable subjects, where no regime change occurs, these four categories are expected to distribute near-uniformly, serving as a negative control.

### Models compared

To test whether greater model capacity rescues post-event prediction, three predictors of very different capacity were trained on the pooled pre-event transitions and evaluated on each subject’s post-event observations: a low-capacity linear baseline (ordinary least squares), logistic regression, and a multilayer perceptron. The perceptron used two hidden layers of 32 and 16 units, ReLU activation, the Adam solver, initial learning rate 0.01, L2 penalty α = 0.001, a maximum of 300 iterations, convergence tolerance 1×10^−3^, and standardized inputs, with a fixed random seed for reproducibility. This comparison asks whether a single model trained on all pre-event data generalizes across a regime-change event.

### Statistical analysis

Between-group comparisons used the two-sided Mann–Whitney U test. CS–DR associations used Spearman rank correlation with 95% bias-corrected bootstrap confidence intervals. Robustness was assessed by the near-uniform distribution of the four categories in stable subjects as a negative control, a 10,000-shuffle subject-level permutation test, propensity matching between the intervention arm and its controls, and leave-one-out iteration over subjects. Analyses used Python (numpy, pandas, scipy, scikit-learn).

### Code availability

All analytic code that computes the cohort construction, phase assignment, per-subject fits, Curvature Shift, Deformation Risk, sign accuracy, the four-category classification, and the model comparison, together with the figures and numerical results, was written by the author in Python and is available from the author on request. Curvature Shift and Deformation Risk are introduced and defined in full above; the companion paper (ref. 6) provides the prior substrate-adequacy framework on which this analysis builds.

## Results

**Table 1.** Cohort composition.

| Arm | Group | n | Min. observations / phase |
| --- | --- | --- | --- |
| Biological-event | Transition subjects | 97 | ≥2 pre, ≥2 post |
| Biological-event | Stable subjects (control) | 212 | ≥2 pre, ≥2 post |
| Interventional | Intervention subjects | 59 | ≥2 pre, ≥2 post |
| Interventional | Untreated controls | 204 | ≥2 pre, ≥2 post |
| Total modeled cohort |  | 309 | — |

Primary biological-event cohort of 309 subjects; the interventional arm and its untreated controls are drawn from the same dataset under identical filters.

**Table 2.** Primary outcomes by regime-change event.

| Quantity | Biological event | Intervention | Stable / control |
| --- | --- | --- | --- |
| Post-event sign accuracy | 0.341 | 0.350 | 0.562 |
| Post-event concordant (%) | 11% | 12% | — |
| Curvature Shift (MMSE/month) | 0.136 (median) | 0.199 (mean) | 0.068 / 0.088 |
| Curvature Shift elevation | 2.23× | 2.26× | — |
| Mann–Whitney P | $4.4 \times 10^{-8}$ | $7.4 \times 10^{-8}$ | — |
| CS–DR coupling (Spearman $\rho$) | 0.500 (95% CI 0.407–0.587) | | — |

Chance sign accuracy = 0.50. The biological-event and intervention columns are structurally independent regime-change events in the same dataset; the stable/control column is the matched negative control.

Of the ADNI participants screened, the primary modeled cohort comprised 309 subjects (97 transition, 212 stable), with a parallel intervention arm of 59 subjects and 204 stable controls. Concordance with the prior trajectory falls to roughly one in nine after either event After the biological event, only 11% of post-event observations remained concordant with prior dynamics; after the intervention, only 12%. In both arms, 62% of post-event readings were silent shift, 11–14% apparent decline, and 13–14% apparent improvement. In stable subjects the four categories were distributed nearly evenly (23% / 28% / 23% / 26%), consistent with noise around a single underlying process and serving as a negative control. At the subject level the pattern was sharper: only 5% in each arm had concordance as their dominant post-event pattern; in the remaining 95%, silent shift or one of the two discordant categories dominated.

### A pre-event predictor is wrong in direction roughly two-thirds of the time

Post-event sign accuracy — the fraction of post-event observations whose direction of change matched the pre-event trajectory — was 0.341 (95% CI, 0.287 to 0.397) after the biological event and 0.350 (95% CI, 0.272 to 0.432) after the intervention, against a chance value of 0.50. The pre-event trajectory assigned the wrong direction to roughly two-thirds of post-event observations, with numerically close failure rates under two structurally independent event types. In stable subjects, post-event sign accuracy was 0.562 (95% CI, 0.528 to 0.596), confirming that the trajectory retained modest directional information where no regime change occurred. The cohort-level deformation of the trajectory law across the event is shown in Figure 2.

**Figure 2.**
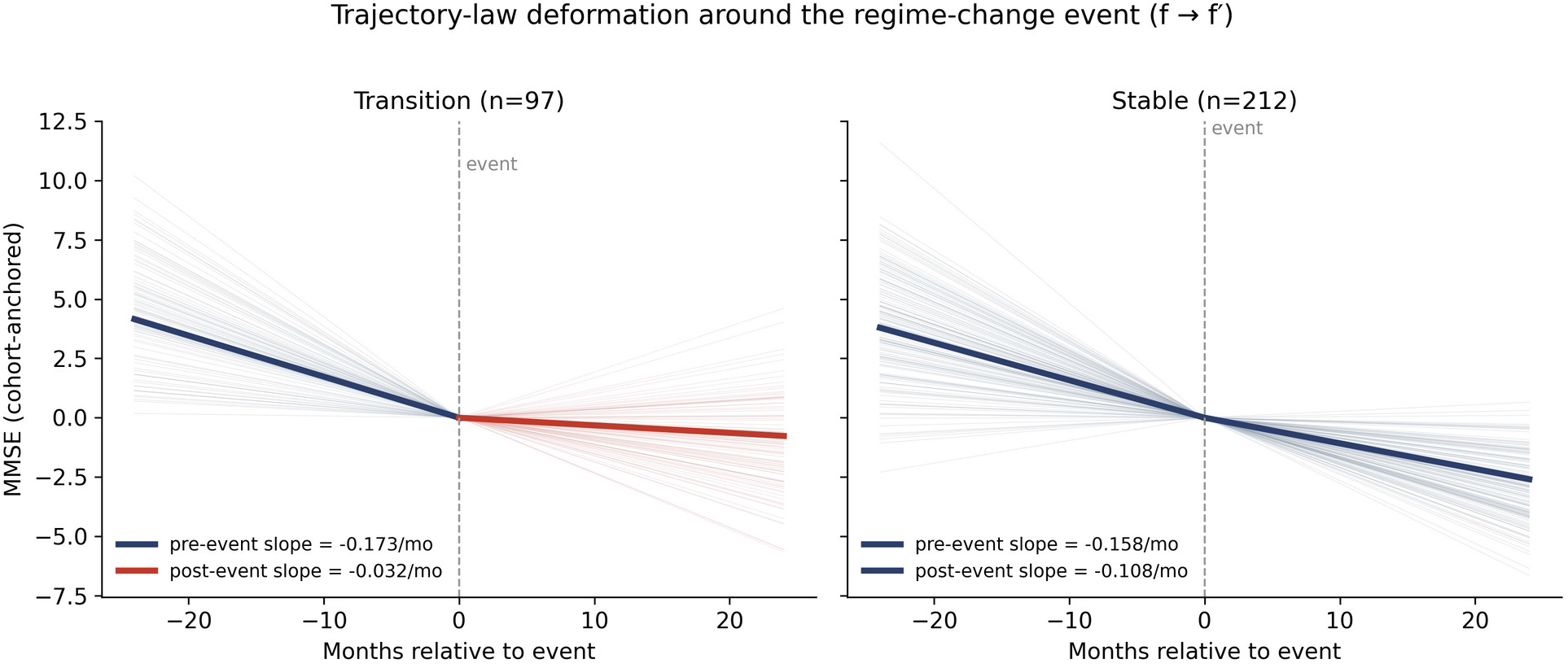
Trajectory-law deformation around the regime-change event (f → f′). Pre- and post-event slopes for transition subjects (n = 97) and stable subjects (n = 212), anchored at the event. In transition subjects the post-event rate departs sharply from the pre-event rate; in stable subjects the rate is approximately preserved.

### Curvature Shift and Deformation Risk quantify the shift and its cost

Median Curvature Shift was 0.136 MMSE/month after the biological event versus 0.068 in stable subjects — a 2.23-fold elevation (Mann–Whitney P = 4.4×10-8). Mean Curvature Shift was 0.199 after the intervention versus 0.088 in controls — a 2.26-fold elevation (P = 7.4×10-8). Deformation Risk inflated 21.9-fold after the biological event and 8.2-fold after the intervention relative to pre-event residuals. Across the full cohort, Curvature Shift and Deformation Risk were coupled (Spearmanρ= 0.500; 95% CI, 0.407 to 0.587; P = 5.8×10-21 ; Figure 3), with tighter coupling within transition subjects (ρ = 0.587) than within stable subjects (ρ= 0.382). The coupling strengthened with deeper post-event for transition subjects (n = 97) and stable subjects (n = 212), anchored at the event. In transition subjects the post-event rate departs sharply from the pre-event rate; in stable subjects the rate is approximately preserved. observation, fromρ= 0.500 at one post-event observation toρ= 0.585 at four. A larger shift in the generating trajectory corresponds to greater failure of the pre-event reference.

**Figure 3.**
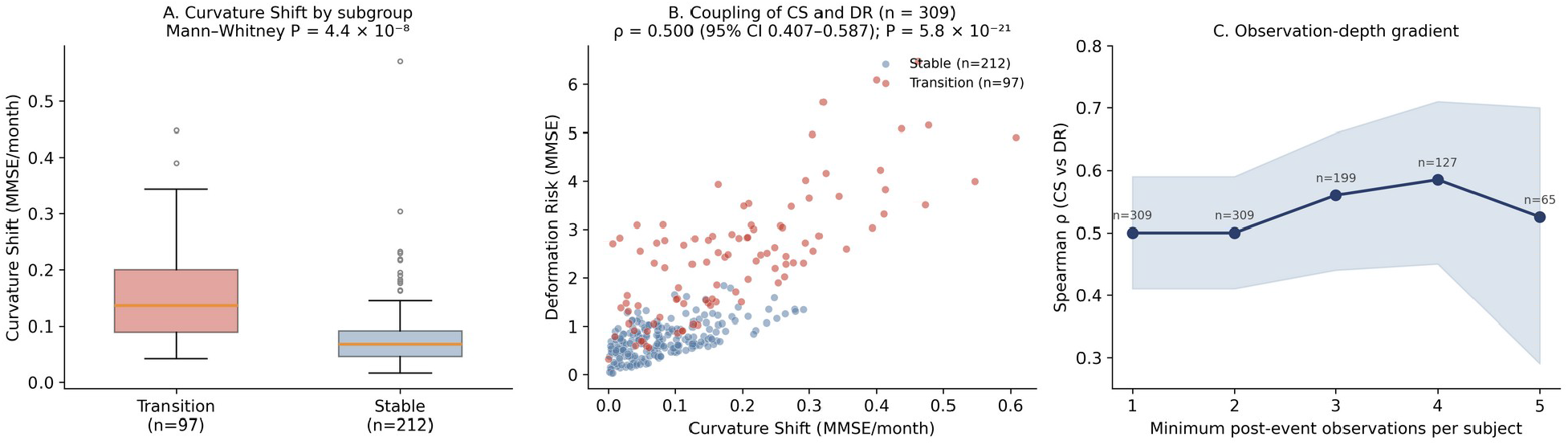
Curvature Shift and Deformation Risk. (A) Curvature Shift is higher in transition subjects than in stable subjects (Mann– Whitney P = 4.4×10^−8^). (B) The two metrics are coupled across the cohort (ρ = 0.500; 95% CI 0.407–0.587; P = 5.8×10^−21^). (C) The coupling strengthens with the minimum number of post-event observations per subject.

### Added model capacity reproduces the failure rather than repairing it

Three predictors of very different capacity — linear, logistic, and a multilayer perceptron — were fit on the pooled pre-event transitions and evaluated on each subject’s post-event observations. Their post-event performance was materially identical: on transition subjects, post-to-pre error ratios were within a few percent of one another, and sign accuracy clustered near 0.30 for all three. The apparent advantage of the higher-capacity model in some strata reflected majority-class prediction in a uniformly declining cohort rather than genuine recovery of direction. Three model classes of very different capacity converging on the same failure pattern show that capacity cannot resolve it.

### The primary signal is robust across subgroups and sensitivity analyses

Post-event sign accuracy in transition subjects remained in the 0.29–0.45 range across demographic strata, and CS–DR coupling remained significant across subgroups of sufficient size. Findings held in a stricter cohort and replicated on the independent endpoint (CS ratio 2.58×;ρ= 0.78). Propensity matching preserved the Curvature Shift elevation at 2.28-fold, addressing confounding between the intervention and its indication. A 10,000-shuffle permutation test produced no permutation more extreme than observed in either arm. Under leave-one-out iteration every headline metric deviated by no more than a few percent. in stable subjects (Mann–Whitney P = 4.4×10-8). (B) The two metrics are coupled across the cohort (ρ= 0.500; 95% CI 0.407–0.587; P = 5.8×10-21). (C) The coupling strengthens with the minimum number of post-event observations per subject.

## Discussion

After a biological transition or the onset of an intervention, the prior trajectory usually no longer governed the subject. The failure pattern was numerically close across two structurally independent events: post-event sign accuracy was 0.341 after the biological event and 0.350 after the intervention — the pre-event reference assigned the wrong direction to roughly two-thirds of post-event observations in both arms — with concordance of 11% and 12% and Curvature Shift elevated 2.23-fold and 2.26-fold relative to their respective controls. Replication on an independent endpoint indicates the signal is not specific to one measurement instrument. Once the generating process changes, interpreting a new observation as a continuation of the prior trajectory becomes unreliable. An effective intervention, by definition, changes how the process evolves; so does a biological transition. The two arms observe the same phenomenon from different causes. The dominant post-event pattern was silent shift. In these observations the direction of change remained consistent with the prior trajectory, but the magnitude and rate did not. This is the most easily missed failure mode, because nothing in a single reading reveals that the subject has moved onto a different trajectory. Apparent worsening and apparent improvement are the more visible misreadings; silent shift is the more common one. The relevant failure is therefore not simply an incorrect point prediction; it is continued interpretation against a law that no longer governs the data. The contribution here is not superior next-observation prediction but the explicit construction of abaselinetrajectoryagainstwhichlaterobservationscanbetested, andasubstrate-levelcriterionfor whether a departure from that baseline carries the geometric signature of an interventional change. Curvature Shift isolates the change in rate that distinguishes a process-modifying intervention from a level shift, and Deformation Risk measures the degree to which the prior law has ceased to govern the subject. Both are computed from the subject’s own longitudinal record, without model weights, counterfactuals, or an assumed causal model — which is what makes the distinction a property of the data substrate rather than of any particular model or causal specification. That higher-capacity models reproduced rather than repaired the pattern indicates that the problem is structural, not a deficit of model flexibility. The two kinds of change cannot always be separated by cause from observation alone; confound- ing between an intervention and the condition that prompts it is, in general, not recoverable from observational data without further assumptions. What the substrate does carry is the geometric signature of a change in the generating law — a change in rate, detectable as Curvature Shift — in- dependent of whether that change was biological or interventional in origin. The metrics determine when a distinction of this kind is supportable from the data and when it is not.

## Limitations

ADNI is a research cohort with higher observation density than many longitudinal datasets, which constrains external validity. The intervention arm was modest, though the main signal persisted after propensity matching. The analysis does not adjudicate the efficacy of any intervention; it identifies when the prior trajectory has ceased to govern observations and whether the departure carries the geometric signature of an interventional change. Measurement-instrument variability would attenuate rather than create directional signal. Replication on an independent endpoint supports the main conclusion.

### Conclusions

In ADNI, only 11–12% of post-event readings remained consistent with the prior trajectory, and a pre-event reference assigned the wrong direction to roughly two-thirds of post-event observations, under both a biological transition and an intervention. The value of AI here is not better point prediction but the explicit construction of the prior trajectory against which later observations are judged, together with a substrate-level criterion — Curvature Shift and Deformation Risk — for whether a departure carries the geometric signature of an interventional change. The metrics identify when that trajectory has stopped governing the subject and under what circumstances the biological-versus-interventional distinction is supportable from the data substrate alone.

## Data Availability

ADNI source data are available to qualified investigators by application, governed by the ADNI Data Use Agreement, which controls all redistribution.

## Use of Artificial Intelligence

A large language model was used to review code for errors and to copy-edit manuscript prose; all scientific content, claims, analyses, and final text are the author’s own and were verified by the author. All analytic code, figures, tables, and numerical results were produced by the author in Python.

